# Evolutionary association of receptor-wide amino acids with G protein coupling selectivity in aminergic GPCRs

**DOI:** 10.1101/2021.09.15.460528

**Authors:** Berkay Selçuk, Ismail Erol, Serdar Durdağı, Ogun Adebali

## Abstract

G protein-coupled receptors (GPCRs) induce signal transduction pathways through coupling to four main subtypes of G proteins (G_s_, G_i_, G_q_, G_12/13_), selectively. However, G protein selective activation mechanisms and residual determinants in GPCRs have remained obscure. Herein, we performed an extensive phylogenetic analysis and identified specifically conserved residues for the receptors having similar coupling profiles in each aminergic receptor. By integrating our methodology of differential evolutionary conservation of G protein-specific amino acids with structural analyses, we identified selective activation networks for G_s_, G_i1_, G_o_, and G_q_. To validate that these networks could determine coupling selectivity we further analyzed Gs specific activation network and associated it with the larger TM6 tilt which is a signature of Gs-coupled receptors. Through molecular dynamics simulations, we showed that previously uncharacterized Glycine at position 7×41 plays an important role in both receptor activation and G_s_ coupling selectivity by inducing a larger TM6 movement. Finally, we gathered our results into a comprehensive model of G protein selectivity called “sequential switches of activation” describing three main molecular switches controlling GPCR activation: ligand binding, G protein selective activation mechanisms and G protein contact.

## Introduction

G protein-coupled receptors (GPCRs) constitute a significant group of membrane-bound receptors that contain five different classes (Fredriksson, Lagerström, Lundin, & Schiöth, 2003; Rosenbaum, Rasmussen, & Kobilka, 2009). The aminergic subfamily of receptors are present in class A and include receptors for dopamine, serotonin, epinephrine, histamine, trace amine, and acetylcholine (Vass et al., 2019). With a large amount of known coupling profiles, experimental structures, and mutagenesis experiments available, aminergic receptors are by far the most studied subfamily of GPCRs. These receptors can couple with different heterotrimeric G proteins which induce distinct downstream signaling pathways (Wettschureck & Offermanns, 2005). Disruption of the proper receptor activation is likely to be the cause of diseases such as coronary heart disease (Jialu Wang, Gareri, & Rockman, 2018) or major depression (Catapano & Manji, 2007; Senese, Rasenick, & Traynor, 2018). Therefore, understanding the molecular mechanisms of coupling selectivity is crucial for developing better therapeutics and diagnostics.

With the advancement of new methodologies, two recent studies have revealed the G protein-coupling profiles of a large set of receptors. Inoue et al. (Inoue et al., 2019) have used a shedding assay-based method to measure chimeric G protein activity for 11 unique chimeric G proteins representing all human subtypes and 148 human GPCRs. Because they have not managed to find an evident conserved motif determining G protein selectivity between receptors, they have built a machine learning-based prediction tool to identify sequence-based important features for each G protein. Similarly, Avet et al. (Avet et al., 2020) have used a BRET-based method detecting the recruitment of the G protein subunits to the receptor to reveal coupling profiles for 100 different receptors. The main strength of this study is that it does not require a modified G protein. Although both high-throughput studies largely agree with each other for certain G proteins, there are inconsistencies between the datasets. Thus, these valuable resources should be analyzed together in detail to gain more power in identifying the selectivitydetermining factors in G protein coupling.

Several attempts have been made to identify molecular determinants of G protein coupling. Most of these (Chung et al., 2011; Du et al., 2019; Liu et al., 2019; Okashah et al., 2019; Semack, Sandhu, Malik, Vaidehi, & Sivaramakrishnan, 2016) have focused on the G protein-coupling interface by analyzing contacts between receptor and the G protein. The others (Kang et al., 2018; Rose et al., 2014; Van Eps et al., 2018; Jinan Wang & Miao, 2019) have highlighted the structural differences between receptors that couple to different G proteins. Flock et al. (Flock et al., 2017) have analyzed the evolutionary conserved positions of orthologous and paralogous G proteins and proposed the “lock and key” model. According to their model, G proteins (locks) have evolved with subtype-specific conserved barcodes that have been recognized by different subfamilies of receptors (keys). Because receptors with distinct evolutionary backgrounds can couple to the same G protein, receptors also must have evolved to recognize the existing barcodes. Although the model has explained the selectivity determining interactions between G protein and receptors, we still lack subfamily specific receptor signaling mechanisms that involves but not limited to the G protein coupling interface.

Despite the extensive research carried out to identify the determinants of G protein selectivity, selectivity determining positions within receptors have remained underexplored. Here, we developed a novel methodology to identify a set of specifically conserved residues for the receptors sharing similar coupling profiles through identification of orthologous receptors. Structural analyses revealed that specifically conserved positions are part of G protein specific activation pathways that allow receptors to transduce signal from ligand binding pocket to the G protein-coupling interface, induce the necessary conformational changes to get coupled by the relevant G protein subtype.

## Results

After a gene duplication event, paralogous clades might diverge from each other with respect to their functions. Therefore, evolutionary pressure against paralogous genes might differ. To perform a precise conservation analysis, we aimed to identify the gene duplication nodes in aminergic receptor evolution. We identified receptor subfamilies (orthologous and paralogous sequences) through a meticulous phylogenetic analysis. As we previously proposed (Adebali, Reznik, Ory, & Zhulin, 2016), the variations that observed in a paralog protein of interest may not be tolerated in the orthologous proteins. In our analyses, we only used orthologous receptors to define a subfamily of interest, members of which are likely to retain the same function. This approach greatly improved the sensitivity of conserved residue assignment for each human GPCR.

To link receptor evolution to its function, we identified residues that are conserved within the functionally-equivalent orthologs for each aminergic receptor. For the residues that play a role in common receptor functions we expect both clades to retain the amino acid residues with similar physicochemical properties. On the other hand, the positions that serve receptor-specific functions, in our case the coupling selectivity, we expect to see differential conservation **(Figure 1a)**. Therefore, we grouped receptors based on their known coupling profiles for eleven different G proteins **(Figure 1b)**. We termed these groups as couplers (e.g., G_s_ coupler receptors) and non-couplers, and performed a two-step enrichment method (Figure 1b) to distinguish specifically conserved residues in couplers from non-couplers. Initially, we used a specific approach to identify evident differentially conserved amino acid residues with high confidence. With the specific approach, residues were labeled as specifically conserved when there was a variation between coupler and non-coupler receptors but not within coupler receptors (Figure 1b. red and blue arrows). This approach depends solely on the coupling profile datasets (Avet et al., 2020; Inoue et al., 2019) and thus, they may contain false-positive couplings. To tolerate the insensitivity introduced by potential false positive couplings, we developed and employed a sensitive approach enabling to obtain a more complete set of residues for each G protein subtype by allowing minor variations within the coupler receptors. With this method, we used a single comprehensive multiple sequence alignment that combined all coupler receptors and their orthologs (Figure 1b. orange arrows), allowed minor variations within a group. We didn’t apply sensitive approach to G_12_ and G_13_ because the low number of coupler receptors would likely cause a high number of false positives. Finally, we compared each aminergic receptor and identified positions that were conserved across all aminergic receptors (consensus) to link the specifically conserved residues to the general mechanism of receptor activation. In total, we identified 53 specifically conserved and 22 consensus residues. The distribution of the specific residues for each G protein is presented in **Figure 1c**.

**Figure 1:**
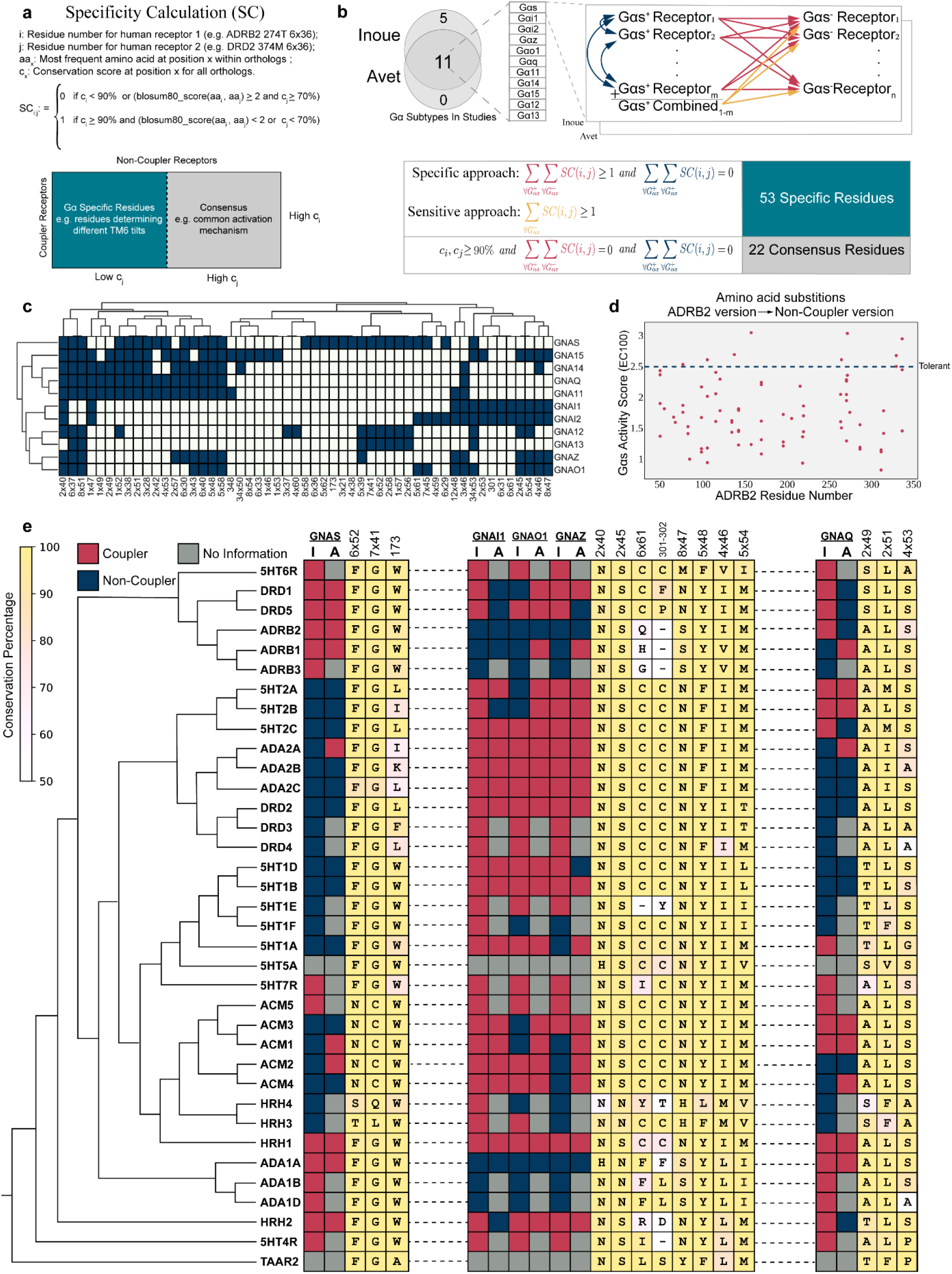
Selectivity determining residues for each Gα subtype. **(a)** The formula for specific residue identification. **(b)** The schema describes the comparisons between paralogous human receptors to find the specifically conserved residues for each Gα. Arrows represent a single comparison. **(c)** The distribution of specifically conserved residues for each Gα subtype and hierarchical clustering of them (complete linkage). **(d)** Possible variants of Gs specific residues that are observed in non-coupler receptors are compared with the wild-type activity score. **(e)** Maximum-likelihood phylogenetic tree of aminergic receptors including coupling profiles, conservation information of selected specifically conserved residues (I: Inoue A: Avet), The background color scale for each consensus amino acids correlates with their conservation (identity).

We aimed to validate the functional impact of potentially deleterious variants that we observe within non-coupler receptors. Thus, we used a dataset (Jones et al., 2020) containing G_s_ activity scores at EC100 for each possible mutation of ADRB2. 31 residues were identified for G_s_ and the activity scores of non-coupler variants were plotted **(Figure 1d)**. Non-coupler variants that we identified predominantly decrease G_s_ activity when compared to average activity of tolerant substitutions. Under normal conditions, the decrease in G_s_ coupling can be attributed to various reasons including misfolding and decreased cell surface expression. However, the substitutions we proposed are not likely to disrupt general receptor functions because the substituting amino acids are indeed found and tolerated in non-coupler receptors (**Figure 1e**) having very high sequence and functional similarity. Additional to the G_s_ coupling dataset, Kim et al. (Kim et al., 2020) mutated two of the residues we identified for G_q_ coupling (8×47 and 6×37) to alanine and showed a decrease in G_q_ activity compared to wild-type 5HT2A receptor which validates that variations at specifically conserved positions are not well-tolerated.

Experimentally shown non-coupler variants cause loss of function in receptors. However, losing the coupling function may not be associated with G protein coupling selectivity. For an amino acid to be involved in G protein coupling selectivity, it should govern functional G protein-specific roles. These roles can be recognition of G protein, ligand binding and/or establishing allosteric receptor conformations that may favor (or disfavor) the engagement with certain G protein subtypes. Hence, we manually assigned each residue into functional clusters such as coupling interface and ligand binding. For example, our method identified positions that are at the G protein coupling interface such as 8×47 (Kim et al., 2020; Maeda, Qu, Robertson, Skiniotis, & Kobilka, 2019; Zhuang, Xu, et al., 2021) and 6×36 (Rasmussen et al., 2011; Xiao et al., 2021; Yang et al., 2020) with no structural information taken into account. The residues that are in the coupling interface are in line with the model that Flock et al. proposed and are likely important for proper G protein recognition. However, for the residues that we could not directly assign a role in G protein coupling activity, we hypothesized that they should be a part of a network controlling the signal transduction from ligand binding pocket to G protein coupling interface and establish required selective structural conformations. To test this hypothesis, we explored the residue-level contact changes upon coupling to a G protein. We used an algorithm that is called Residue-Residue Contact Score (RRCS) which has been proposed to identify the common activation mechanism in class A GPCRs (Zhou et al., 2019). We calculated ΔRRCS for each interacting residue pairs by subtracting contact scores of the active structure from the inactive structure. All the active structures we used contained a heteromeric G protein machinery coupled to receptor. We filtered out residue pairs with |ΔRRCS| <= 0.2 and only kept residues that are in our pool of conserved residues (75 residues in total). We analyzed structures of eight different receptors with four different G proteins (see Methods). The structures we used were experimentally characterized except for one state of a single receptor. As we aimed to use the 10 active-state G_αs_ coupled structures of DRD1, which lacks an experimental inactive structure, we used a model inactive DRD1 structure (Pándy-Szekeres et al., 2018) retrieved from GPCRdb (Kooistra et al., 2021).

In total, we analyzed 41 pairs of active and inactive structures and identified ΔRRCS values of activation networks. We analyzed each network and detected edges (increase or decrease in contact score) observed at least 36 times regardless the sign of ΔRRCS value to build a network that would represent all 41 networks. By using this network, we identified the most frequently used signal transduction paths **(Figure 2a)**, connecting ligand binding pocket to G protein-coupling interface and create a basis for the routes that can induce coupling selectivity. We divided the receptor into five layers based on sequential nature of interactions and illustrated the direction of signal transduction between layers. Additional to the 4 layers (1-4) that were previously proposed in the common activation mechanism (Zhou et al., 2019) we defined “Layer 0” which is corresponds to the ligand binding site. Though the most of the signaling paths pass through important motifs such as Na^+^ binding pocket and PIF (Katritch et al., 2014), it is remarkable that the novel path starting with a 3×37 does not require the involvement of any of these important motifs. Within the identified network, the signal is transmitted from ligand binding pocket to the G protein interface by using mainly TM2, TM3, and TM4. We projected all the residues onto an inactive structure of ADRB2 based on the layers they belong to (**Figure 2b**) to provide an insight about their locations of different layers.

**Figure 2:**
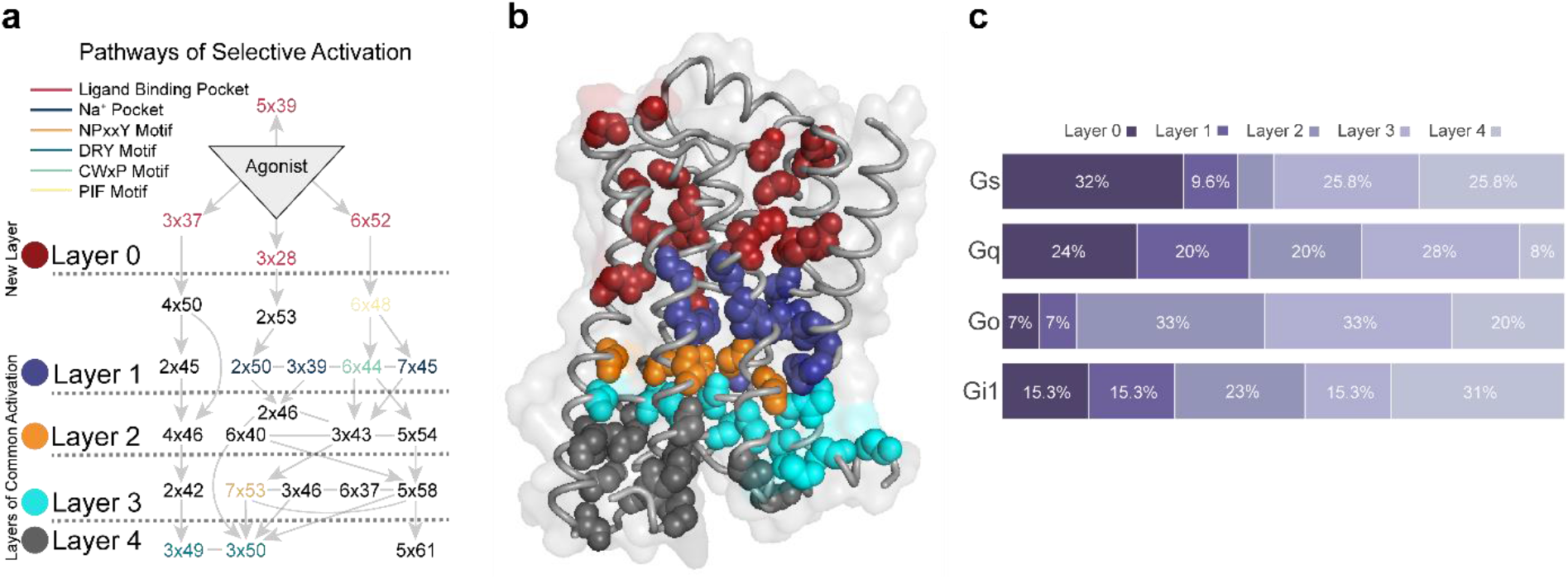
Structural analysis of molecular pathways that are observed upon coupling with heteromeric G protein complex. **(a)** The most common molecular signal transduction pathways from ligand binding pocket to G protein coupling interface. The arrows represent a contact change upon coupling to a G protein. The network is summarized and divided into different layers based on their functional relevance. **(b)** Projection of main chains of specifically conserved and consensus residues in different layers of activation on inactive ADRB2 structure (PDB ID 2RH1) **(c)** The distribution of specifically conserved residues for each analyzed Gα subtype.

To determine the contribution of each layer for G_s_, G_i1_, G_o_ and G_q_, we calculated the distribution of specific residues to different layers **(Figure 2c)**. Layer 0 and Layer 1 are more involved in the coupling for G_s_ and G_q_ relative to G_i1_ and G_o_. For G_o_, 86% of the coupling-related residues are positioned in the layers (2, 3 and 4) closer to the G protein binding site. Differences in these distributions indicate mechanistic differences between distinct coupling events.

To detect if the specifically conserved residues have differential roles in G protein coupling-related mechanisms, we grouped ΔRRCSs (contact changes upon coupling to a G protein) for the receptors coupled to same G protein and compared with the rest by using two sample t-test. This approach yielded interaction changes (ΔΔRRCS) within the receptors that are significantly different (p<0.01) and specific for G_s_, G_i1_, G_o_, and G_q_. Significant contact changes occurring between 75 conserved residues were used to construct G protein specific activation mechanisms. The constructed networks (**Figure 3b-e**) support our evolution-driven hypothesis and demonstrate that specifically conserved residues indeed have differential mechanistic roles in G protein coupling. In parallel to the Figure 2c, networks for Gs and G_q_ contained ligand contacting residues (Figure 3a and Figure 3e) while networks for Gi1 and Go do not. Although, Gi1 and Go belong to same subfamily and they share 8 of the specifically conserved residues (47% of the specifically conserved residues for G_o_ and 62% for G_i1_) of G proteins their networks are totally different from each other. Moreover, even when we grouped the receptors coupled to Gi together, no significant difference in contact scores having p-value less than 0.01 was observed (Supplementary Table) for the shared specifically conserved residues (Figure 1c). This suggests that receptors coupling to Gi may not necessarily share a common activation mechanism. Therefore, these differences in activation networks could be one of the factors determining selectivity between G_i1_ and G_o_ coupled receptors.

**Figure 3:**
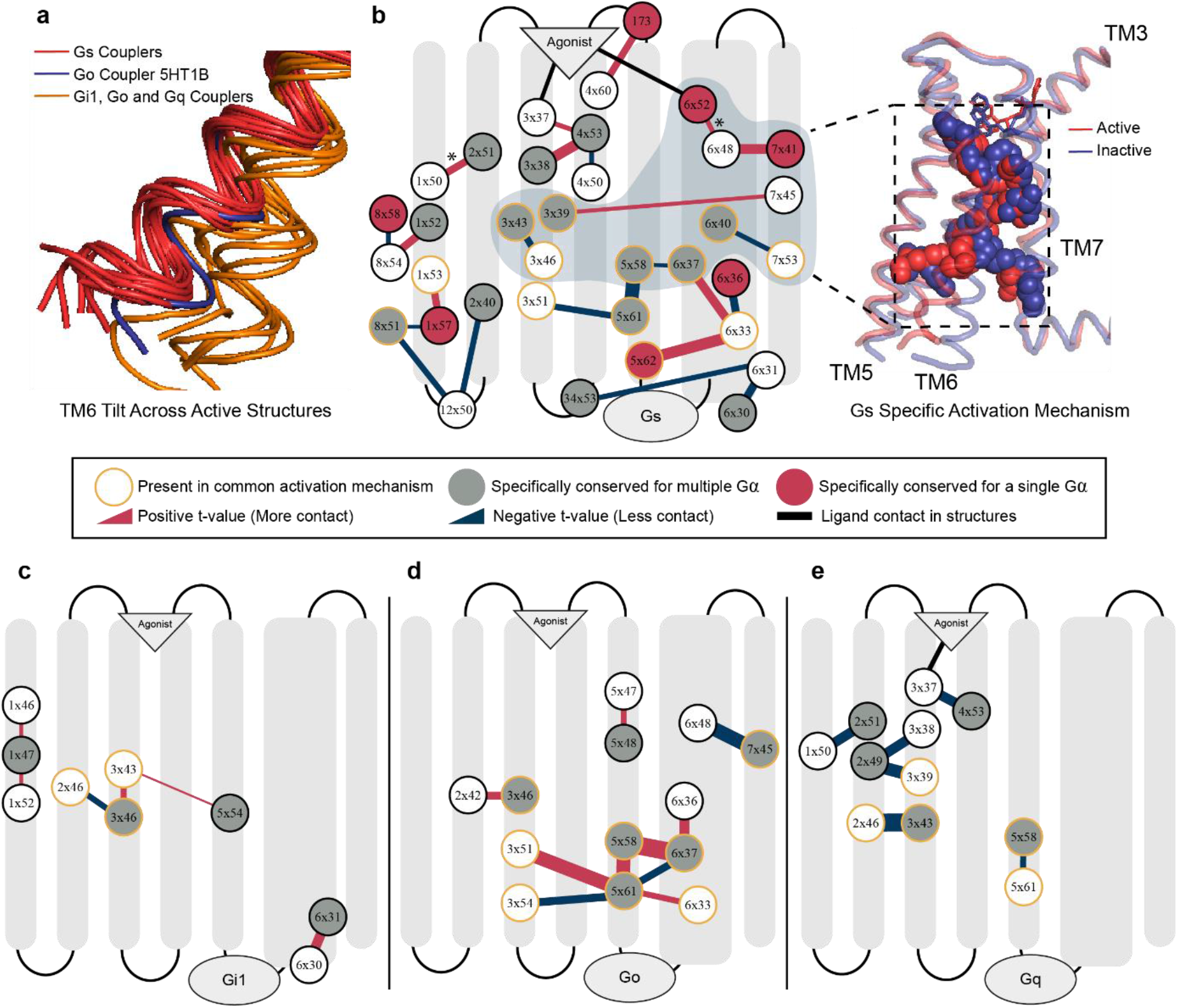
Specific Activation Networks for G_s_, G_i1_, G_o_ and G_q_. **(a)** TM6 tilt comparison between the active receptors we used. Red: G_s_ couplers, Orange: G_o_, G_i1_ and G_q_, Blue:5HT1B G_o_ coupler as an exception. **(b)** Interactions within the receptor that are specific (p<0.01) to G_s_. Red: increasing contact, blue: decreasing contact, orange circle: present in common activation mechanism, red fill: uniquely identified specific residue for G_s_, grey fill: G_α_ specific residue. Width of the lines correlate with statistical significance. Group of residues that possibly facilitate in TM6 movement for G_s_ coupling were shown on inactive (blue) and active (red) structures. (**c-e)** Specific interaction networks for G_i1_, G_o_ and G_q_. p<0.1 is used for G_i1_. *: This interaction is identified only if 5HT1B is neglected from the comparison due to its larger TM6 movement.

Even though residues specifically conserved for the receptors sharing similar coupling profiles are part of G protein specific activation networks, it is still not clear that these contact changes are the basis for selective coupling, or they arise due to the physical interaction with a G protein itself. To show that these networks can determine selectivity we further analyzed the activation network for G_s_ coupled receptors. Previously, it was shown that receptors coupled to G_s_ achieve a larger TM6 tilt (Rose et al., 2014; Van Eps et al., 2018) than the receptors coupled to other G proteins. Superimposition of the active structures that we used in our analysis **(Figure 3a)** is also in line with the previous findings. We hypothesized that the network we identified can modulate this structural difference. Furthermore, requirement for a larger TM6 movement can be the reason why G_s_ specific activation mechanism is more complex than the rest (Figure 3b-e). An exception to this is the TM6 position of 5HT1B (García-Nafría, Nehmé, Edwards, & Tate, 2018) that is coupled to G_o_ (Figure 3a, blue structure), because it achieved a slightly larger tilt. Thus, we performed an additional statistical test to reveal possible interactions that can promote larger TM6 movement by excluding the samples for 5HT1B and revealed the 6×52-6×48 interaction indicating the role of 6×48 in differential TM6 movement in G_s_ coupled receptors. (p=0.0023).

We projected a part of G_s_ specific activation network which we predicted to be associated with the differential TM6 movement onto experimentally resolved active (red, 3SN6) and inactive (blue, 2RH1) ADRB2 structures **(Figure 3b)**. More specifically, we hypothesized that the network containing 6×52 and 7×41 triggers this structural difference because interactions at the upper layers are more likely to be leading a structural change. In agreement with our hypothesis, deep mutational scanning of ADRB2 (Jones et al., 2020), has revealed that 7×41 is the second and 6×48 is the fourth most intolerant residue to any mutations and, to our knowledge, no previous study has identified the functional role of 7×41 until now. It is expected that a position that is crucial for G_s_ coupling to be to be one of the most intolerant residues for a receptor primarily coupled to G_s_.

To validate our methodology and further understand the mechanistic insight of the relevance of core transmembrane region in G protein coupling, we studied the glycine at position 7×41 as a test case and performed molecular dynamics (MD) simulations. We applied three different mutations, G315C, G315Q, and G315L, on monomeric active and inactive-state ADRB2 **(Figure 4a)**. We particularly selected variants observed in acetylcholine and histamine receptors (Figure 1e) to validate our hypothesis that variants in non-coupler aminergic receptors at the same position are inactivating. We used two main metrics to assess the molecular impact of these three mutations. First, the comparison active/inactive states based on GPCRdb distances (see methods) revealed that wild-type receptor keeps its active state more than the variants **(Figure 4b)** and leucine residue was the most inactivating mutation. The significant inactivation through integration of leucine mutation is parallel to pre-existing experiments (Arakawa et al., 2011; Jones et al., 2020). Then, to identify the molecular changes in absence of glycine, we evaluated the significant contact differences (ΔRRCS) between WT and mutated MD simulation trajectories.

**Figure 4:**
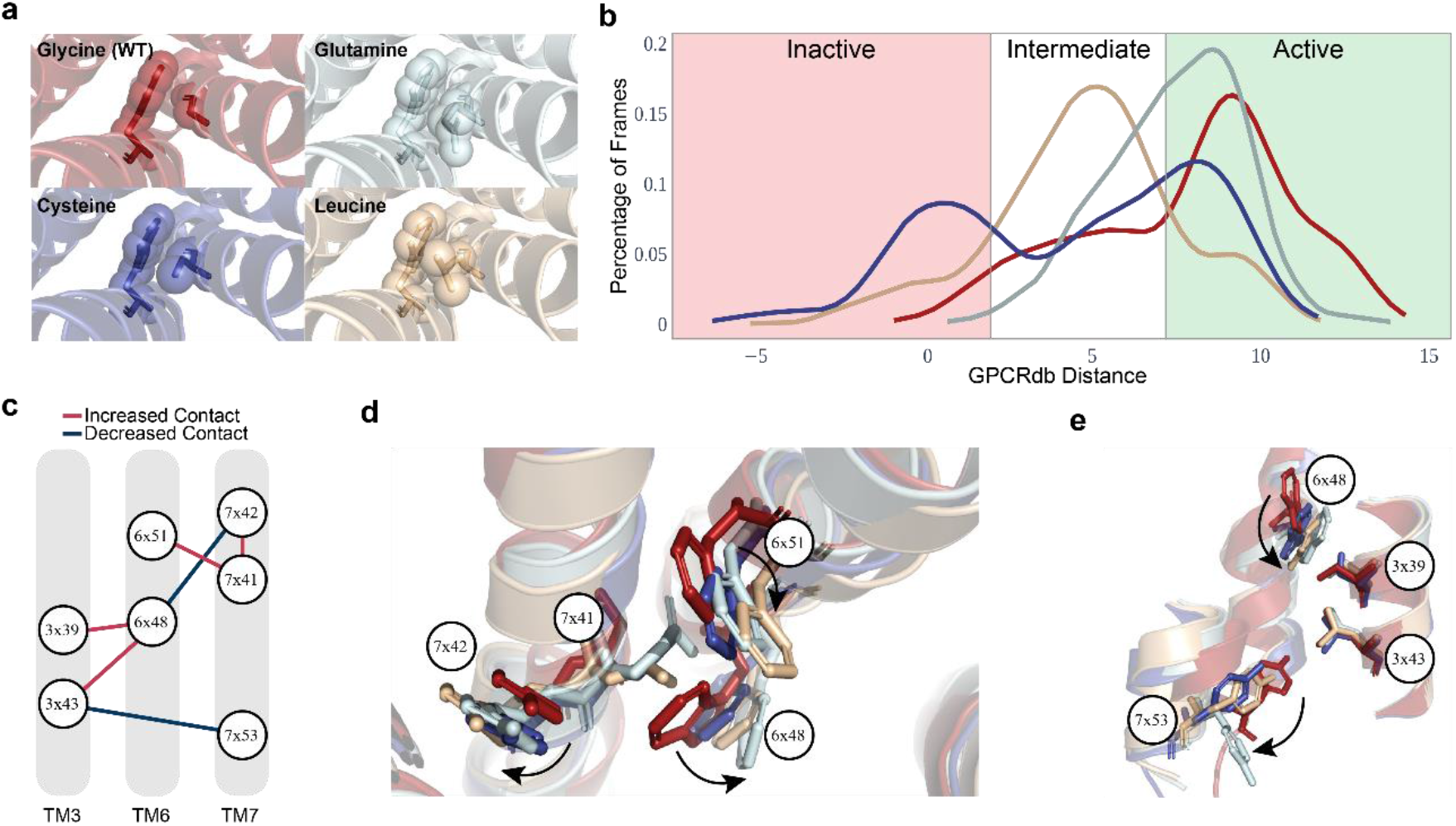
Analysis of molecular dynamics simulations reveal functional importance of glycine at 7×41. **(a)** 4 different MD simulation systems were shown in their initial conformation. **(b)** For each simulation distribution of frames with respect to their state of activation were shown, distance in Angstrom. **(c)** The common pathway representing impact of the mutations at 7×41. **(d-e)** The common pathway was represented on average structures that were obtained in all MD trajectories for every mutation and WT. The movements of residues were represented with arrows.

To examine the entire trajectory, we selected 11 frames from each simulation with 50 ns time intervals (in total 500 ns) for each replicate. Thus, we compared residue-residue contact scores of 77 mutated and 77 WT frames for active-state simulations, while we compared 22 mutated and 22 WT frames for inactive-state simulations by using two-sided t-test. For each mutation and activation state, we identified significant contact changes (p<0.01) and intersected common changes that we observed for all of mutated systems. As a result, we identified 135 residue pairs for active and 83 residue pairs for inactive simulations. When we projected these residue pairs (135 residue pairs) as a contact network, we identified a conserved and highly affected pathway **(Figure 4c)** connecting ligand binding pocket to NPxxY motif which showed changes towards inactivation of the receptor. Then, we projected the identified molecular pathway onto average cluster structures that were produced by using the trajectories from all 7 replicates (35000 frames in total) for each mutation **(Figure 4d-e)**. MD results suggested a pathway (Figure 4c) which explains the importance of G315: An increase bulkiness of the amino acid at 7×41 (by non-glycine amino acids) leads to increased contact with 7×42 and 6×51 while 7×41 physically impairs the interaction between 6×48 and 7×42. When 6×48 loses its contact with 7×42 (Figure 4d), it increases its contact residues at TM3 3×43 and 3×39 (Figure 4e). Increased interactions between TM6 and TM3 loosens TM3-TM7 packing which is an important initiator of the TM6 tilt in class-A GPCRs (Zhou et al., 2019). Additionally, it loosens the contacts between TM6 and TM7 through 6×48-7×42, 6×44-7×49, and 6×52-7×45, which explains the increased distance between 7×53 and 3×43 (Figure 4e). Moreover, the simulations of cysteine and leucine variants exhibited an increased contact between 3×43 and 6×40 (p<0.01) inhibiting the receptor activation through restricting outward TM6 movement. When we evaluated the inactive trajectories, we observed similar contact changes between 6×48, 6×51, 7×41 and 7×42 (p<0.01) proving that the simulation results are not biased to active-state simulations. Thus, analysis of MD trajectories suggests that glycine at 7×41 plays an important role in receptor activation, and it is likely to control selectivity for G_s_ coupling by promoting a larger tilt of TM6 which we observe almost exclusively in Gs coupled receptors.

## Discussion

By integrating our findings and current literature we propose a G protein selectivity model involving a series of modules. As pilots turn on switches in a pre-determined order before the takeoff, GPCRs must turn on their molecular switches for a specific type of G-protein coupling to occur. If pilots fail to turn on all the switches properly due to an error, there will be no permission for them to depart. Similarly, all molecular switches must be turned on for receptors to engage with a G protein and induce downstream signaling pathways. For these reasons, we named our model **“sequential switches of activation”.** We propose the existence of three main switches within a GPCR structure. The first switch checks for binding of the proper agonist which induces conformational changes in lower layers of the receptors. If an agonist makes the proper contacts with the receptor the first switch turns on. Then as a next step, receptors should be activated through G protein selective activation mechanisms which includes multiple micro-switches to turn of the second main switch. Micro-switches represent the arrangement of inner contacts that are specific for G protein subtypes. When inner contacts are established properly the second switch turns on as well. As a third and last check point, receptors should contain the set of residues that can recognize the ridges on G proteins according to the “key and lock” model that Flock et al suggested. When required contact between G protein and receptor is established, the third switch turns on and the receptor is successfully coupled by a subtype of G proteins. Mutations inducing constitutional activity can be considered as a “short circuit” because they can bypass switches. On the other hand, mutations that halt receptor’s ability to turn on a particular switch can prevent coupling. It is important to note that our model is inclusive of and complementary to the model Flock et al. suggested. Combination of these two models gives us a more complete perspective on receptor-level determinants of coupling selectivity.

**Figure 5:**
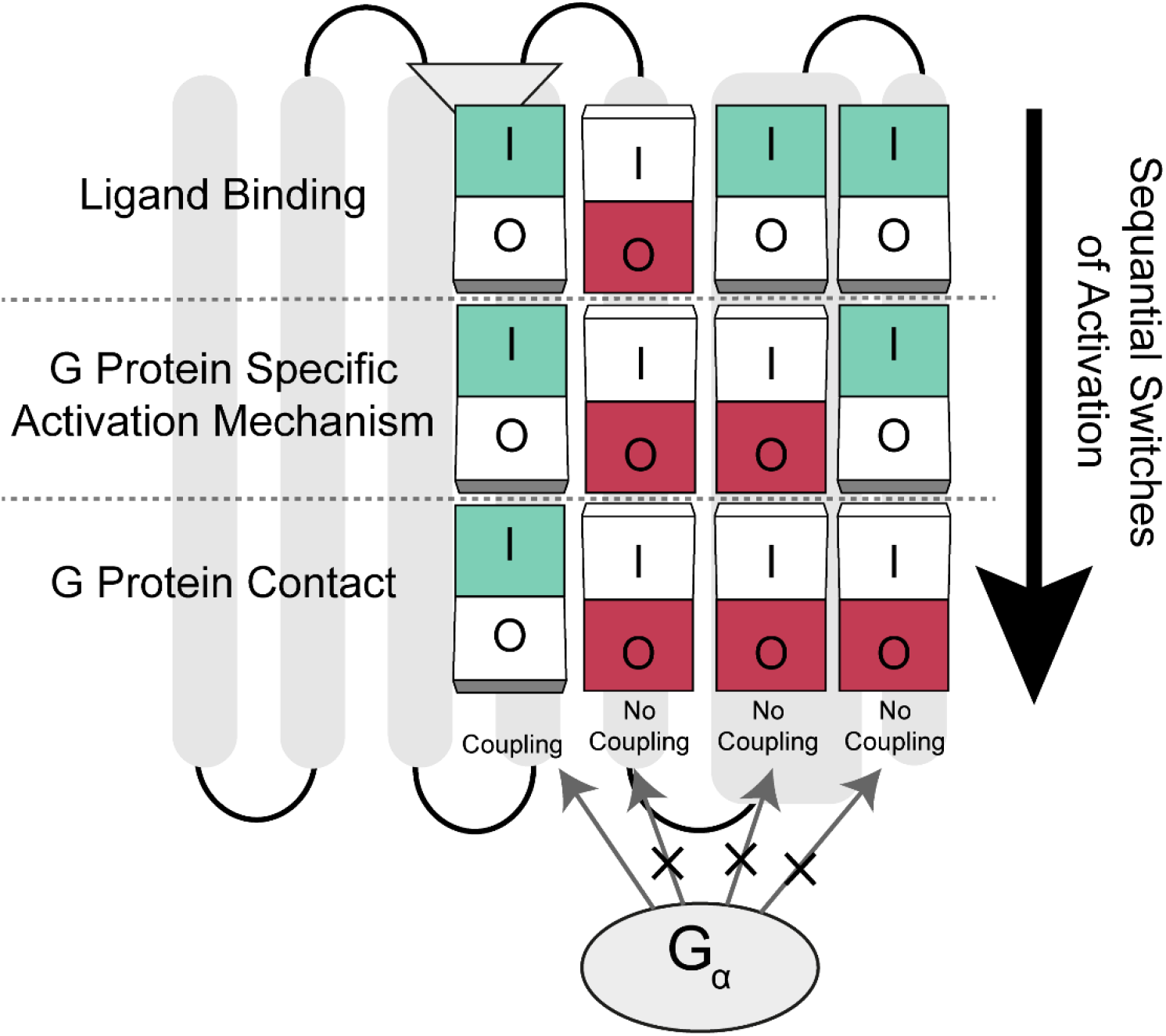
Sequential switches of activation model for G protein selectivity. The model describes that all switches in different layers of receptors must be turned off for receptor activation and coupling of the G protein. If switches at upper layers are halted due to a mutation, following switches become turned off which inhibits G protein coupling eventually.

In our study, we used a novel phylogenetic approach to identify residues that are conserved among groups of receptors coupling to a particular G protein. We identified the largest possible set of residues (Figure 1c) by combining sensitive and specific approaches together. Due to our greedy approach while some positions could determine coupling selectivity, others may be “passenger” positions that may modulate core receptor functions. Moreover, the positions we identified are the ones that are shared among all aminergic receptors and lack receptor-level variations. Previous studies on chimeric GPCRs (Wess, 1998, 2021; Wong, 2003) point out the importance of ICL3 in determining coupling selectivity. While we identified residues that contact with G proteins, our analyses did not reveal any possible determinants at ICL3. This indicates that the determinants at ICL3 are not shared between aminergic receptors and rather be specific to individual receptors. Alternatively, in nature there may not be a solution for G protein coupling selectivity determination with ICL3 only. Experimentally constructed chimeric receptor activation should be handled with caution because they cannot be evaluated as a part of receptor evolution. Thus, to identify all selectivity-determining positions, each receptor should be analyzed individually.

Although our study does not include any direct experimental evidence that coupler or non-coupler variants alter coupling selectivity, it provides sufficient evidence to support the existence of receptor-wide selectivity determinants not only at the G protein coupling site but throughout receptors including the ligand binding site. We used G_s_ coupling data from deep mutational scanning of ADRB2 performed by Jones et al. to show that non-coupler variants cause loss of function (Figure 1d) (Jones et al., 2020), their roles in determining coupling selectivity should be clarified further. With that purpose, we used residue-residue contact score algorithm and revealed involvement of specifically conserved residues in G protein specific activation mechanisms (Figure 3b-d) which suggests their role in determining coupling selectivity. We should note that due to scarcity of G_q_, G_o_ and G_i1_ coupled structures, the networks we provided could be modified in the future as the number of G protein coupled experimental structures increase. As a third layer of evidence, we identified the role of a previously uncharacterized G^7×41^ (Jones et al., 2020) for ADRB2 and G_s_ coupled receptors through molecular dynamics simulations (Figure 4c).

Although we cannot rule out the potential effect of G^7×41^ in non-G_s_ activation, we can conclude that it has a critical importance for determining G_s_ coupling selectivity. The fact that G^7×41^ is dispensable for Gi couplers suggest that it may not be as critical for those GPCRs and Gi activation. To summarize, multiple layers of evidence suggest that G protein selectivity determinants for aminergic receptors are distributed receptor-wide.

The conclusions of this study are limited aminergic receptors only because there has been no supporting evidence for a common selective mechanism that might present for all class A GPCRs. Therefore, it is necessary to handle each GPCR subfamily separately to identify subfamily specific selectivity determinants. With such an effort, it may be possible to discover commonalities and differences between different subfamilies of GPCRs. Although different subfamilies of receptors couple to a G protein by having similar structural conformations, underlying mechanisms for achieving a conformation might vary. As the number of solved G protein-coupled receptor structures increase in the protein data bank, it is inevitable that new selectivity determinants and similar mechanisms will be discovered in near future.

## Acknowledgements

This work is supported by EMBO Installation Grant (to OA 4163) that is funded by TUBITAK. The molecular dynamics simulations reported in this paper were fully performed at TUBITAK ULAKBIM, High Performance and Grid Computing Center (TRUBA resources).

## Methods

### Sequence Selection

Sequence selection is the very first step of this study. We used the BLAST+ (Camacho et al., 2009) algorithm to obtain homologous protein sequences from other organisms. We blasted a human target protein to find its homologs. The UniProt (“UniProt: a worldwide hub of protein knowledge,” 2019) database is used as a source for the sequences. We retrieved all the sequences until the third human protein from the blast output.

### Multiple Sequence Alignment (MSA) #1

After sequence selection, the next step is performing multiple sequence alignment for obtained sequences. For this purpose, we used MAFFT (Katoh & Standley, 2013) “einsi” option which allows large gaps. This option allows us to align multiple homologous regions of different receptors.

### Maximum Likelihood (ML) Tree #1

The MSA was used to produce a maximum likelihood (ML) tree. ML trees helped us to find relationships between different proteins. ML Tree 1 was used to identify the clade which contains our protein of interest. For ML tree construction we use the IQ-Tree version 2.0.6 (Minh et al., 2020) We used 1000 Ultra-fast bootstraps and JTT+I+G4+F substitution model. IQ-Tree is used at this step for mainly its high speed in bootstrapping.

### Obtaining Gene Clade

For making modifications on the ML trees we use a Python based tool ETE3 (Huerta-Cepas, Serra, & Bork, 2016). To analyze a tree, we first need to root it properly. We chose the third human protein from our BLAST results, as an outgroup. Then, we traversed from our target human leaf node to root until we reached a clade containing another human protein. After each move, we analyzed the species content of the clades we are observing. When a clade contained species that were not observed in previous moves, we included all of the leaf nodes to our analysis. On the other hand, when a clade contains a previously observed species, we exclude that clade from our analysis, because seeing a species at a lower phylogenetic levels is an indication of a differential gene loss event. We continued with the remaining sequences and produced a multiple sequence alignment with them.

### Multiple Sequence Alignment Trimming

MSA trimming is needed to remove some of the noise from the alignment and it speeds up tree reconstruction. MSA trimming removes positions that are misleading for tree production. For example, positions having too many gaps can be removed from the alignment. We used trimAl (Capella-Gutiérrez, Silla-Martínez, & Gabaldón, 2009) with automated1 option which is stated to be the best option for constructing maximum likelihood trees.

### Maximum Likelihood Tree #2

ML tree 2 was used to identify the paralogous sequences that we have in our analysis. For ML tree construction we used the RaxML-NG version 0.9.0 (Kozlov, Darriba, Flouri, Morel, & Stamatakis, 2019) --search option with JTT+I+G4+F substitution model.

### Paralog Trimming

Paralog trimming is a key part of our approach. After gene duplication, one of the paralogous clades tends to diverge more than the other. Unless the diverged clade is removed from our analyses (MSA), it might introduce false divergence signals in conservation calculation. For this reason, we need to exclude diverged paralogs from our analyses. We used the second ML tree for detection of the diverged paralogs.

We first calculated the global alignment scores (BLOSUM62 is used) of every sequence on the ML tree 2 with respect to our human target sequence. We assessed each internode having two child clades based on the number of leaf nodes and species they contain. When two child clades contained at least one identical species, we looked for significant divergence between the clades in terms of global alignment scores to label one clade as paralogous. Also, we need those clades to be evolutionarily comparable, thus we compared the taxonomic level of the organisms between two clades. If the clades are comparable to each other, we applied two-sample t test for by using the global alignment scores. If one clade has significantly lower similarity scores (p<=0.1) that clade is labeled as a diverged paralogous clade. We applied the same approach for detecting the taxonomic level of the organisms and common lineage numbers with *Homo sapiens* was used this time (p<=0.1). If the clades are evolutionarily comparable and one clade had a significantly lower global alignment score, all of the sequences belonging to that clade were eliminated.

When two of the clades contained less than three sequences each, it was hard to obtain a significance. Therefore, for those cases we compared the average global alignment scores and eliminated the clade with lower average. For the remaining situations we don’t remove any of the clades.

### Conservation Calculation

After obtaining orthologs we used them to calculate the conservation scores for each receptor.

The conservation percentage for a certain residue is calculated as follows:

1. Find the most frequent amino acid for a certain position in the multiple sequence alignment (MSA).
2. After finding the most frequent amino acid, we compared it with other alternatives in that position. When comparing amino acids, we calculated BLOSUM80 score for each of them. If the BLOSUM80 score is higher than 2 we accept it as an “allowed” substitution because it means that these amino acids replace each other frequently and have similar properties.
3. The gaps are not included while calculating the conservation percentage.
4. If gaps are more than %50 percent, we categorized that position as a gap.
5. The conservation score is equal to the number of most frequently observed and “allowed” amino acids over number of all non-gap positions

### Specificity Calculation (SC)

For a position to be specific or consensus the criteria is the following:

1. First, we need one alignment of two proteins with their orthologs. Then we split the alignment into two alignments with the same length.
2. We label a position as **consensus**, when both alignments are conserved more than consensus threshold (90%) at that particular position and the most frequent amino acids are similar (BLOSUM80 score is more than 1) to each other.
3. We calculated conservation percentages for each alignment. There are two different scenarios in this case. The first one is when the most frequent amino acids of the two of the alignments are not similar (BLOSUM80 score is lower than 2) to each other. If this is the case and conservation percentage for any alignment is above the specificity threshold (90%) we label that position as **specifically conserved** for that alignment. The second case is where the most frequently observed amino acids are similar to each other. In this case, for a position to be specific for one alignment first it should satisfy the specificity threshold and secondly the conservation percentage of the other alignment should be lower than our lower threshold (70%).

For the steps above we choose 90 percent for both specificity and consensus thresholds. 70 percent is selected for lower specificity threshold.

### Enrichment of Specifically Conserved Residues

We identified specifically conserved residues with two different approaches:

#### Specific Approach

1. We divided receptors into two as couplers vs non-couplers. Let’s assume that we have n number of couplers and m number of non-couplers.
2. We compare coupler receptors with non-couplers in a pairwise manner. In these comparisons we count the number of being specific for every residue. In total there are n times m comparisons. We divide the obtained counts to the total number of comparisons in order to get the frequency of a residue being specific for the couplers’ group.
3. To examine if a residue is generally variable or specific to the coupling event, we compared couplers with themselves. We applied STEP 2 for couplers-couplers comparison as well. This time, we have n*(n-1) comparisons in total. We again calculated the frequencies accordingly.
4. For the specific approach, we don’t allow any inside variation and this makes the result of STEP 3 zero. On the other hand, for a residue to be labeled as specific, we expect the STEP 2 more than zero. When these two conditions are satisfied, we label that residue as specifically conserved

#### Sensitive Approach

1. We built a comprehensive multiple sequence alignment for the coupler receptors and their orthologs.
2. We compared this alignment with non-coupler receptor’s MSAs similarly to the STEP 2 of the Specific Approach.
3. We added newly discovered positions to our analysis as specifically conserved.

#### Building the maximum-likelihood phylogenetic tree for aminergic receptors

1. We blasted (Camacho et al., 2009) aminergic receptors and obtained first 50 sequences to generate a fasta file.
2. From that fasta file we selected representative sequences by using cd-hit default options.
3. MAFFT (Katoh & Standley, 2013) einsi algorithm was used to align representative sequences.
4. IQTree version 2.0.5 (Minh et al., 2020) was used to create the phylogenetic tree with options: -m JTT+G+I+F -b 100 --tbe

### Residue-Residue Contact Score (RRCS) and Network Analysis

We calculated the RRCS score for 20 active (ADRB2: 3SN6,7DHI; DRD1: 7CKW, 7CKX, 7CKZ, 7CKY, 7CRH, 7JV5, 7JVP, 7JVQ, 7LJC, 7LJD; DRD2: 6VMS, 7JVR; DRD3: 7CMU, 7CMV; 5HT1B: 6G79; ACM2: 6OIK; 5HT2A: 6WHA; HRH1: 7DFL)(García-Nafría et al., 2018; Kim et al., 2020; Maeda et al., 2019; Rasmussen et al., 2011; Xia et al., 2021; Xiao et al., 2021; Xu et al., 2021; Yang et al., 2020; J. Yin et al., 2020; Zhuang, Krumm, et al., 2021; Zhuang, Xu, et al., 2021) and 24 inactive structures (ADRB2: 2RH1, 6PS2, 6PS3, 5D5A; DRD1: GPCRdb inactive model; DRD2: 6CM4, 6LUQ, 7DFP; DRD3: 3PBL; 5HT1B: 4IAQ, 4IAR, 5V54, 7C61; ACM2: 3UON, 5YC8, 5ZK3, 5ZKB, 5ZKC; 5HT2A: 6A93, 6A94, 6WH4, 6WGT; H RH1: 3RZE)(Cherezov et al., 2007; Chien et al., 2010; Fan et al., 2020; Haga et al., 2012; C.-Y. Huang et al., 2016; Im et al., 2020; Ishchenko et al., 2019; Kim et al., 2020; Kimura et al., 2019; Miyagi et al., 2020; Shimamura et al., 2011; Suno et al., 2018; C. Wang et al., 2013; S. Wang et al., 2018; W. Yin et al., 2018). For each receptor we substracted inactive RRCS from activeRRCS to obtain ΔRRCS values for each residue pairs. We wrote a custom python code to obtain files with ΔRRCS scores. We combined all of the networks that contain information about the contact changes upon activation to produce the most common molecular signal transduction pathways. (Supplementary File). For the details of the RRCS algorithm please read the corresponding article (Zhou et al., 2019).

### Identification of G protein Specific Activation Networks

After obtaining ΔRRCS networks for each active-inactive structure pairs we grouped ΔRRCS values based on the G protein subtype coupling the receptors. Then we compared ΔRRCS values of individual groups (e.g. G_s_: ADRB2 and DRD1) with the rest of the groups (e.g. Non-G_s_: DRD2, DRD3, 5HT1B, ACM2, 5HT2A, HRH1) by using two-sample t-test. While p<=0.01 is used for G_s_, G_q_, and G_o_, p<=0.1 is used for G_i1_. We obtained significant contact changes upon coupling to a particular G protein.

### Molecular Dynamics Simulations

We downloaded inactive and active structures of Beta2 Adrenergic receptor (β2AR) from PDB (PDB ID: 4GBR, and 3SN6, respectively)(Rasmussen et al., 2011; Zou, Weis, & Kobilka, 2012). Three thermostabilizing mutations, T96M^2×66^, T98M^23×49^, and E187N^ECL2,^ were mutated back to the wild-type (WT) in both sequences. Since used inactive structure of the β2AR has a short ICL3 that links the TM5 and TM6, we did not introduce additional residues to the ICL3, and used the crystal structure as it is. However, active structure of the β2AR lacks ICL3, and we modeled a short loop with GalaxyLoop code (Park, Lee, Heo, & Seok, 2014). We inserted FHVSKF between ARG239 and CYS265. We introduced three changes at 315^7×41^ position, and one WT and obtained three mutants (namely; G315C, G315L, and G315Q). We used PyMOL to place mutations (PyMOL(^™^) Molecular Graphics System, Version 2.1.0.). Orientations of proteins in biological membranes were calculated with OPM server (Lomize, Pogozheva, Joo, Mosberg, & Lomize, 2012) and We used CHARMM-GUI web server (Jo, Kim, Iyer, & Im, 2008; Lee et al., 2016; Wu et al., 2014) to create input files for the molecular dynamics simulations for Gromacs. Since, inactive and active structures start with ASP29^1×28^ and GLU30^1×30^; end with LEU342^Cterm^ and CYS341^8×59^, respectively, we introduced acetylated N-terminus and methylamidated C-terminus to the N and C-terminal ends. Two disulfide bridges between CYS106^3×25^-CYS191^ECL2^, and CYS184^ECL2^-CYS190^ECL2^ were introduced. Each lipid leaflet contains 92 (1-palmitoyl-2-oleoyl-sn-glycero-3-phosphocholine) POPC biological lipid type (total 192 POPC molecules in system). Systems were neutralized with 0.15 M NaCl ions (50 Na^+^ and 55 Cl^-^ ions in total). We used TIP3P water model for water molecules (MacKerell et al., 1998), and CHARMM36m force field for the protein, lipids and ions (J. Huang et al., 2017). One minimization and six equilibration steps were applied to the systems, before production runs (for the equilibration phases 5 ns, 5 ns, 10 ns, 10 ns, 10 ns, and 10 ns MD simulations were run, in total 50 ns). In equilibration phases, both Berendsen thermostat and barostat were used (Berendsen, Postma, Van Gunsteren, Dinola, & Haak, 1984). In production runs, we applied Noose-Hoover thermostat (Hoover, 1986; Nosé & Klein, 1983) and Parrinello-Rahman barostat (Parrinello & Rahman, 1980). 500 ns production simulations were run with Gromacs v2020 (Abraham et al., 2015) and repeated 7 times to increase sampling (in total for each system we simulated 3.5 μs). 5000 frames collected for each run, and for instance for the WT system, we concatenated 35000 frames to calculate GPCRdb distance distributions (*gmx distance* tool was utilized for this purpose) and find average structures (Visual Molecular Dynamics code utilized to find average structure (Humphrey, Dalke, & Schulten, 1996)). To calculate the GPCRdb distance in Class A GPCR structures, CYS125^3×44^-ILE325^7×52^ distance subtracted from TYR70^2×41^-GLY276^6×38^ distance. If calculated distance is higher than 7.15 Å, lower than 2 Å, and between 2-7.15 Å state of the receptors labelled as active, inactive, and intermediate, respectively (Isberg et al., 2015; Shahraki et al., 2021). All figures were generated with PyMOL v2.1.0. To estimate water accessibilities to the internal cavity of the receptors, and sodium ion accessibilities to the ASP79^2×50^, we calculated averaged water and sodium ion densities. Time averaged three-dimensional water and sodium ion density maps were calculated with GROmaρs (Briones, Blau, Kutzner, de Groot, & Aponte-Santamaría, 2019)

### Analysis of Contact Changes Within Molecular Dynamics Simulation Trajectories

Frames of MD simulation trajectories were selected from 0ns to 500ns with 50ns gaps for each trajectory and replicate for a mutation. Including the frame at t=0ns, for a replicate we obtained 11 frames to represent the whole trajectory. We have applied the same strategy for all 7 active-state replicates and obtained 77 frames for WT and mutated MD trajectories. For each frame, we calculated RRCSs for every residue pair and identified statistically significant (p<0.05) differences between WT and mutated trajectories by applying a two-sided t-test. For the inactive simulations, we had only two replicates, therefore we compared 22 mutated frames with 22 WT frames.

After applying t-test, we intersected the significant contact changes we observed for each mutational state to observe the common change due to the absence of glycine. In total, we identified 135 common changes for active-state simulations and 83 common changes for inactive-state simulations. We used Cytoscape (Shannon et al., 2003) to visualize the changes as a network. PyMOL was used to visualize the identified pathway on protein structures.

## Data and Materials Availability

The open-source code and supplementary data are available at our GitHub repository: https://github.com/CompGenomeLab/GPCR-coupling-selectivity

The MD trajectories are available at: https://doi.org/10.5281/zenodo.5763490.

